# Host adaptations in the *fixLJ* pathway of the *Burkholderia cepacia* complex increase virulence

**DOI:** 10.1101/2020.05.04.070367

**Authors:** Matthew M. Schaefers, Benjamin X. Wang, Nicole M. Boisvert, Sarah J. Martini, Sarah L. Bonney, Christopher W. Marshall, Michael T. Laub, Vaughn S. Cooper, Gregory P. Priebe

## Abstract

The *Burkholderia cepacia* complex (BCC) is composed of multiple species, including *B. multivorans* and *B. dolosa,* that are significant pathogens for people with cystic fibrosis (CF) and are extensively resistant to many antibiotics. The *fixL* gene of the *fixLJ* 2-component system (TCS) in these BCC species shows evidence of positive selection for nonsynonymous mutations during chronic lung infection in CF. Previous work showed that the *B. dolosa fixLJ* system regulates 11% of the genome and modulates biofilm formation, motility, persistence within macrophages, and virulence in a murine pneumonia model. Here, we assess the impacts of clinically observed FixL evolved variants in *fixLJ* pathway-mediated phenotypes in *B. dolosa* and *B. multivorans.* BCC carrying the ancestral *fixL* sequence are less pathogenic than constructs carrying evolved variants in both a macrophage infection model and a murine pneumonia model. *In vitro* phospho-transfer experiments demonstrate that the evolved *B. dolosa* FixL variants are able to reduce *fixLJ* pathway activity by either having lower levels of kinase activity or increased phosphatase activity. Notably, the ancestral *fixL* genotype has increased ability to survive within the soil compared to isogenic constructs with evolved *fixL* genotypes, demonstrating that increased virulence comes at an expense. Modulation of the FixLJ system has profound effects on many BCC phenotypes including full pathogenicity, and this modulation is critical for BCC adaptation to the host.

## Main

The *Burkholderia cepacia* complex (BCC) is a group of more than 20 species of closely-related Gram-negative bacilli that can be dangerous respiratory pathogens for people with cystic fibrosis (CF).^1,2^ Among CF patients in the US colonized with BCC, the species most commonly seen are *B. cenocepacia* and *B. multivorans*, although there is significant variability based on geographic region and institution,^3–7^ and *B. multivorans* has emerged as the most predominant BCC species infecting CF patients in some regions. ^4–7^ BCC members have caused several outbreaks within the CF community,^2^ including one outbreak of a highly antibiotic-resistant strain of *B. dolosa* among almost 40 CF patients in Boston^8^ and another of *B. cenocepacia* in Toronto.^9^ BCC can also cause serious infections in individuals with chronic granulomatous disease (CGD).^10^ Outbreaks of hospital-acquired BCC infections in non-CF and non-CGD patients have also been increasingly described, often associated with contaminated medications,^11–16^ including a recent outbreak associated with contamination of the stool softener docusate with *B. contaminans.*^15,16^

Analysis of genomic diversity arising during *B. dolosa* and *B. multivorans* chronic infections in CF identified the *fixLJ* two-component system (TCS) as a pathway that is under positive selection, evident from enrichment for non-synonymous mutations as opposed to synonymous mutations and genetic parallelism among many independent infections.^17–19^ TCSs are one mechanism that bacteria use to sense and respond to their environment.^20^ Our previous work determined that the *fixLJ* pathway senses oxygen tension, is important for virulence in a murine model of pneumonia, and regulates ~11% of the genome.^21^ Additionally we found that the *fixLJ* system is involved in biofilm formation and motility as *B. dolosa* lacking *fixLJ* made more biofilm and had reduced motility.^21^ The *fixLJ* system was also critical for survival within THP-1-dervived human macrophages. These experiments were conducted using a *fixLJ* deletion mutant to determine the effects of deletion of both genes. In the current study, we found, that constructs carrying clinically observed FixL variants (single amino acid changes) were more virulent, had altered gene expression, and were less able to survive within soil that isogenic constructs carrying ancestral FixL variants. These results highlight the importance of the FixLJ pathway in BCC pathogenesis and provide insight into the evolution of BCC during chronic infection.

## Results

### Mutations in BCC FixL sensory domain are associated with decline in lung function

In a series of published works, involving more than 100 *B. dolosa* longitudinal isolates taken over 16 years from 14 different CF patients and 22 *B. multivorans* isolates taken from a single CF patient over 20 years, we identified mutations within *fixLJ* during chronic lung infection.^17–19^ The locations of these mutations within the predicted domains of FixL are depicted in Figure 1A. Most mutations are within the predicted PAS and PAC domains, which are conserved sensory domains.^22^ Based on our previous work^21^ and the predicted heme-binding pocket, it appears that the BCC *fixLJ* pathway is an oxygen-sensing mechanism. Of note, the FixL proteins of *B. dolosa* (AK34_969) and *B. multivorans* (BMD20_10585) share 98% protein identity.

**Figure 1.**
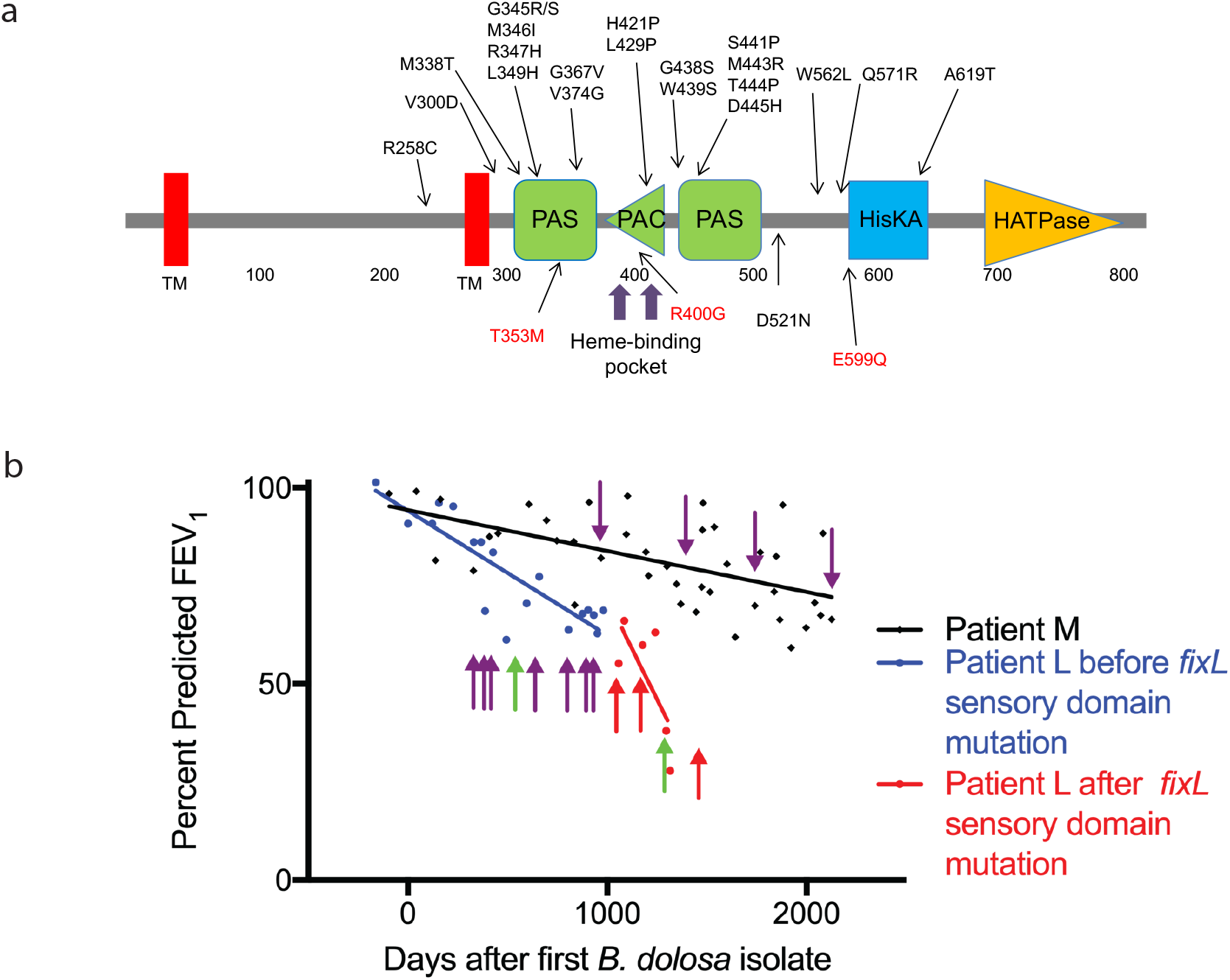
Mutations in the *Burkholderia dolosa* FixL sensory domain are associated with decline in lung function in cystic fibrosis. **a.** Domains predicted by SMART ^65^ and observed SNPs in *B. dolosa* in black ^17,18^ and *B. multivorans* in red.^19^ Domain abbreviations: TM, transmembrane; PAC, Motif C-terminal to PAS motif; HisKA, histidine kinase; HATPase, histidine kinase-associated ATPase. **b**. Rapid decline in percent predicted FEV1 (ppFEV1) in a CF patient (Patient L) after detection of mutations in the sensory domain (PAS) of FixL. Downward arrows denote sequenced isolates from patient M, upward arrows denote sequenced isolates from patient L. Purple arrow: ancestral fixL allele; red arrow: *fixL* mutation in sensory domain; green arrow: *fixL* mutation not in sensory domain. P=0.05 by linear regression, comparing ppFEV1 slopes before and after detection of *fixL* mutations in sensory domain (red arrows).

We compared the lung function of patients from our previous study^18^ who were infected with *B. dolosa* isolates containing FixL sequence variants to determine if there was a correlation between FixL sequence variants and clinical outcomes. An increased rate of decline in lung function, measured by percent predicted FEV1 (ppFEV1), in patient L after *B. dolosa* isolates with mutations in the predicted sensory domain (red arrows) was detected. This slope differs in comparison to earlier time points, before such mutations were detected (Figure 1B, purple and green arrows, red versus blue line: P=0.05 comparing slopes using linear regression). The single green arrow among the purple arrows demonstrates that multiple lineages coexisted at the same time in the lung^17^. Another patient, patient M, who lacked isolates mutated in the predicted FixL sensory domain, had a more modest decline in lung function over a similar period.

To determine the specific phenotypes of evolved FixL alleles, we generated isogenic constructs in reference isolate *B. dolosa* AU1058. The *fixLJ* deletion strain^21^ was complemented with the ancestral FixL sequence (W439) or the evolved sequence found in strain AU0158 (W439S). It is worth pointing out that the reference strain AU0158 is an evolved strain as it was isolated ~3 years after initial *B. dolosa* infection. Strains containing the FixL sequence variants that were associated with lower lung function in Figure 1B were also generated: G345S and R347H, and both constructs had the ancestral tryptophan, W, at residue 439. These constructs were also complemented with the conserved ancestral sequence of FixJ, along with ~600 base pairs upstream allowing for expression from its native promoter, using a mini-Tn7 based vector allowing for long-term stability without the need for selection by the insertion into the chromosome.^23^ We also compared the effects that FixL sequence variants have on virulence in *B. multivorans* by generating isogenic variants within *B. multivorans* VC7102 (BM2) by replacing a fragment of the *fixL* gene in its native location with a fragment with the desired evolved mutation.^24^

### BCC strains carrying evolved FixL variants are more virulent in human macrophages and a murine pneumonia model but less able to survive in soil

We compared the ability of BCC constructs carrying FixL sequence variants to survive within the THP-1 human monocyte cell line treated with PMA to differentiate into macrophage-like cells. *B. dolosa* constructs carrying the evolved FixL variant (W439S) and the evolved variants associated with a period of clinical decline (G345S and R347H) were significantly better able to survive at 2-3-fold higher levels compared to constructs carrying ancestral variant (W439) or lacking *fixLJ* (empty vector) (Figure 2A). One of the two *B. multivorans* evolved variants, FixL T353M, had increased survival within THP-1-dervived macrophages compared to the ancestral strain (VC7102) (Figure 2B).

**Figure 2.**
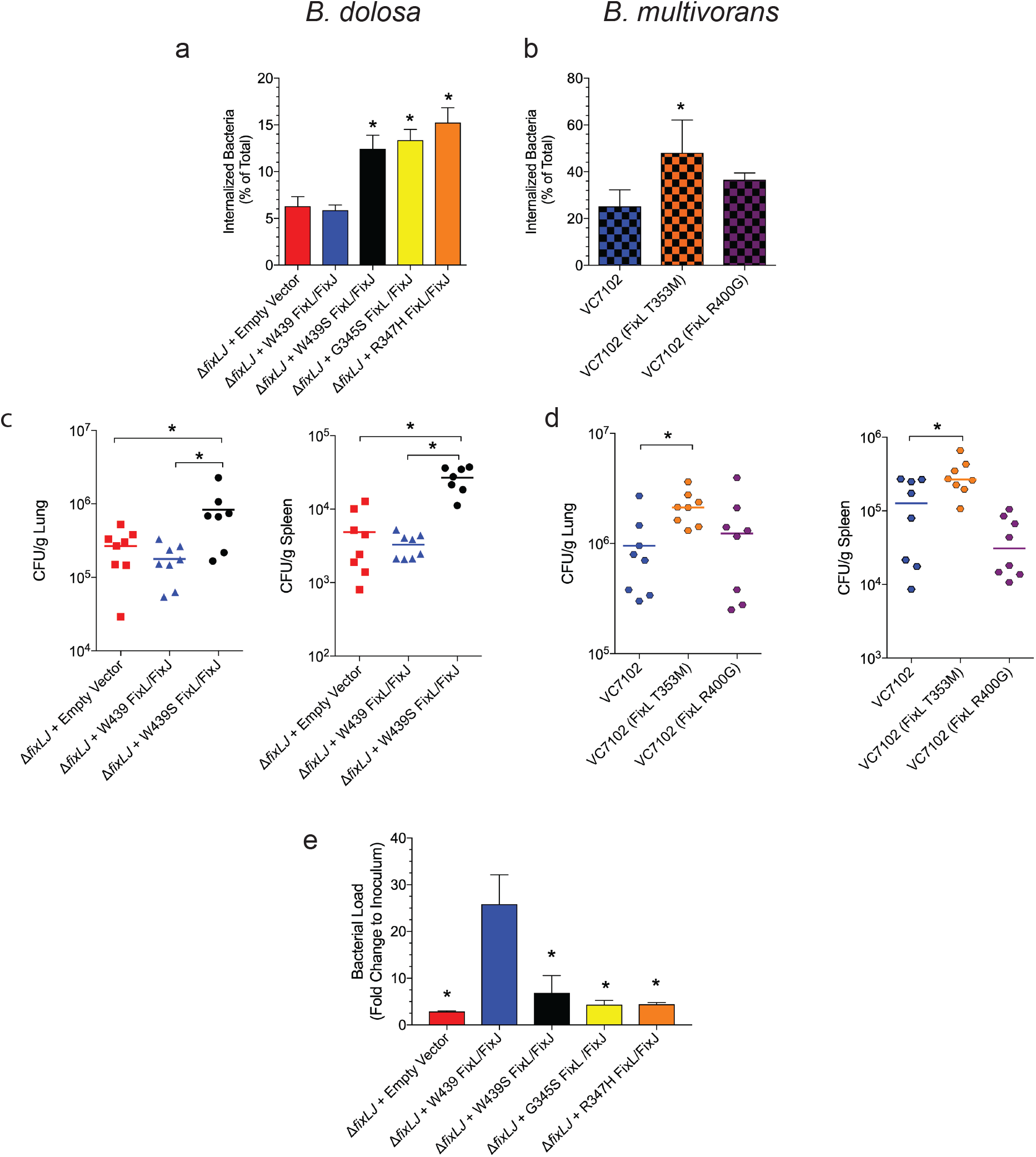
*Burkholderia cepacia* complex strains carrying evolved FixL variants are more virulent in human macrophages and a murine pneumonia model but less able to survive within soil. PMA-treated THP-1 human macrophages were infected with ~2×106 CFU/well (MOI of ~10:1) of **(a)** *B. dolosa* strain AU0158 or **(b)** *B. multivorans* strain VC7102 isogenic constructs carrying FixL sequence variants for 2 hours, after which the percent of internalized bacterial relative to the total bacterial growth was determined by killing extracellular bacteria with kanamycin (1 mg/mL). Means from 2-3 separate experiments with three replicates per experiment are plotted with error bars representing one standard deviation. *P<0.05 by ANOVA with Tukey’s multiple comparison test compared to **(a)** Δ*fixLJ* + empty vector and Δ*fixLJ* +W439 FixL/FixJ or **(b)** strain VC7102. C57BL/6 mice were intranasally challenged with ~4×10^8^ CFU/mouse of **(c)** *B. dolosa* strain AU0158 or **(d)** *B. multivorans* strain VC7102 isogenic constructs carrying FixL sequence variants. Bacterial loads were measured in the lungs and spleen 7 days after infection. Data are representative of 2 separate experiments with 7-8 mice per group. Each point represents one mouse, and bars represent medians. *P<0.05 by ANOVA with Tukey’s multiple comparison test. **(e)** *B. dolosa* strain AU0158 Δ*fixLJ* mutant complemented with *fixLJ* isogenic constructs or empty vector were inoculated into 1 g sterile soil (1-6 ×10^6^ CFU in minimal media) and incubated for 10 days. Bacterial load was measured and plotted relative to the inoculum used. *P<0.05 compared to W439 by ANOVA with Tukey’s multiple comparison test.

Our previous work found a correlation between the ability to survive within THP-1-dervived macrophages and the ability to persist within murine lungs and to disseminate and persist with the spleen after intranasal inoculation.^21^ In order to measure the ability of the constructs to persist *in vivo*, C57BL/6 mice were intranasally infected with these BCC constructs using similar methods. Mice infected with *B. dolosa* carrying the evolved (reference) FixL variant (W439S) had 4-5-fold higher levels of bacteria within the lungs and spleen compared to mice infected with *B. dolosa* carrying the ancestral FixL variant (W439) or lacking *fixLJ* (empty vector) (Figure 2C). Mice that were infected with *B. multivorans* VC7102 carrying the evolved FixL variant that conferred increased survival within THP-1-dervived macrophages (T353M) had significantly increased bacterial loads with the lungs and the spleen compared to mice infected with bacteria carrying the ancestral sequence variant or the other evolved variant not associated with increased survival in THP-1-dervived macrophages (R400G) (Figure 2D).

To determine if the *fixL* mutations that confer an increased level of virulence within the host came with an expense in ability to survive within soil, a natural ecological niche of BCC, we measured the ability of the different constructs to survive within soil for 10 days. We found that all three evolved FixL variants (W439S, G345S, and R347H) had a 5-6-fold reduction in the ability to survive within soil compared to the ancestral variant (W439) (Figure 2E). Bacteria lacking *fixLJ* were also less able to survive within the soil, demonstrating the importance of the *fix* pathway to survival both within the host and in the environment.

### BCC strains carrying evolved FixL variants are more motile, make less biofilm, and have altered gene expression

Our previous work showed that *B. dolosa* lacking *fixLJ* produced increased levels of biofilm and had decreased motility.^21^ We found that both *B. dolosa* and *B. multivorans* carrying evolved FixL alleles produced significantly less biofilm than isogenic strains carrying the ancestral FixL sequence (Figure 3A and 3B). *B. dolosa* and *B. multivorans* constructs carrying evolved FixL sequence variants also had increased motility compared constructs carrying the ancestral variant (Figure 3C and 3D). Interestingly, *B. dolosa* carrying the ancestral FixL variant was completely non-motile, the diameter plotted in Figure 3C (~12 mm) was the diameter of the 10 μL drop placed on the agar surface. *B. dolosa* carrying an empty vector which lacks *fixLJ* still had the ability to swim albeit at lower levels (Figure 3C).

**Figure 3.**
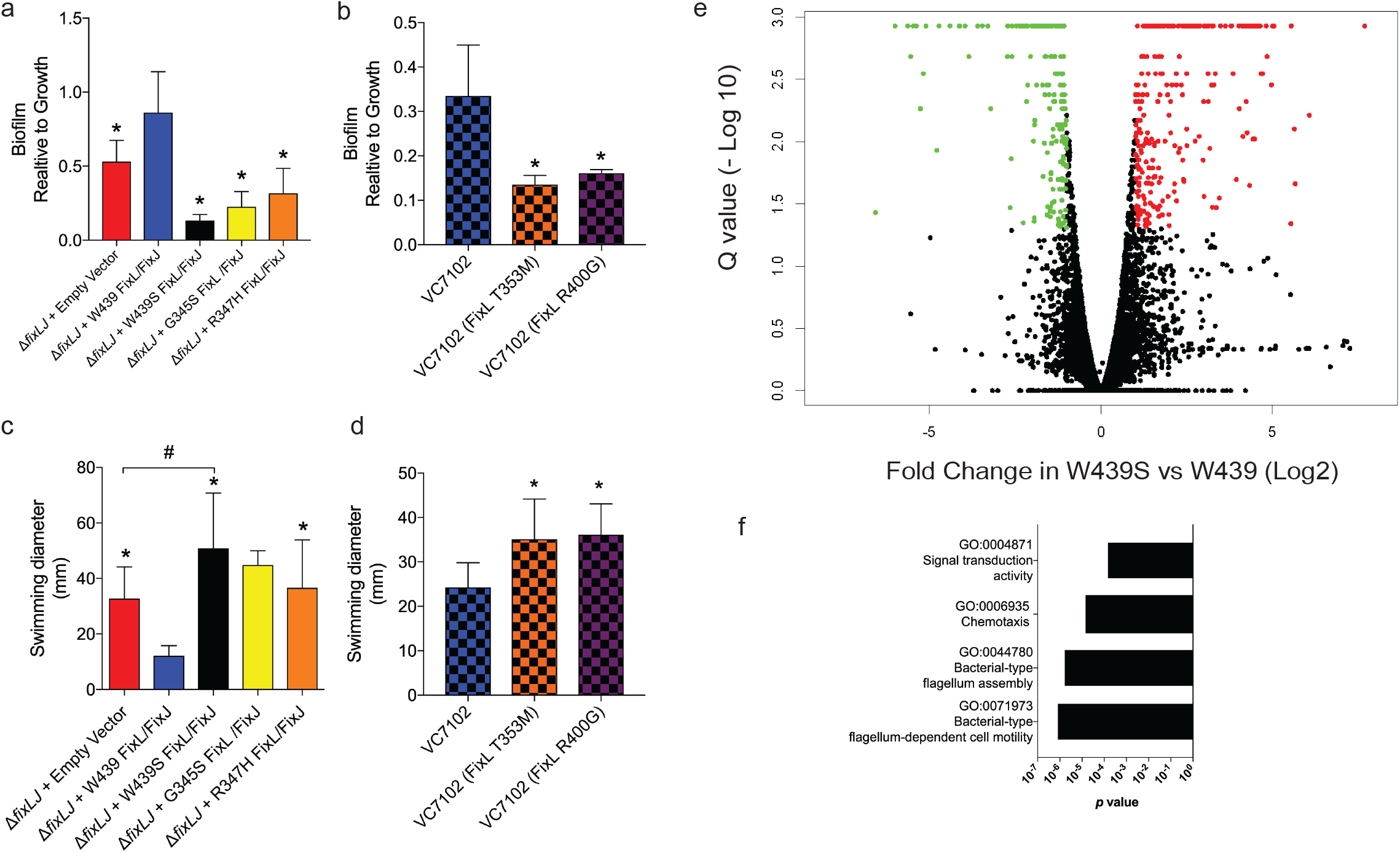
*Burkholderia cepacia* complex strains carrying evolved FixL variants are more motile, make less biofilm, and have altered gene expression. Biofilm formation of **(a)** *B. dolosa* strain AU0158 or **(b)** *B. multivorans* strain VC7102 isogenic constructs carrying FixL sequence variants on PVC plates as measured by crystal violet staining at 48 hours. Means from 3 separate experiments with 5-6 replicates per experiment are plotted with error bars representing one standard deviation. *P<0.05 by ANOVA with Tukey’s multiple comparison test to Δ*fixLJ*+W439 FixL/ FixJ or VC7102. Motility of **(C)** *B. dolosa* strain AU0158 or **(D)** *B. multivorans* strain VC7102 isogenic constructs carrying FixL sequence variants on low-density (0.3%) LB agar and swimming distance was measured after incubation for 48 hours. Means from 3 separate experiments with 3-4 replicates per experiment are plotted with error bars representing one standard deviation. *P<0.05 by ANOVA with Tukey’s multiple comparison test compared to construct carrying ancestral FixL variant (W439 or VC7102); #P<0.05 by ANOVA with Tukey’s multiple comparison test. **(E)** Volcano plot depicting the differential regulation of genes. Green dots signify genes with expression 2-fold lower in the *B. dolosa* strain AU0158 construct carrying the ancestral FixL variant (W439) relative to a construct carrying the evolved variant (W439S), with a q < 0.05. Red dots signify genes with expression 2-fold higher in the evolved variant (W439S) construct carrying the relative to a construct carrying the ancestral FixL variant (W439), with a q < 0.05. **(F)** GO terms that were enriched with an adjusted p value <0.05 among genes that were statistically upregulated (q value <0.05, at least 2-fold) in *B. dolosa* carrying the evolved FixL variant (W439S) relative to *B. dolosa* carrying the ancestral FixL variant.

To identify the differentially expressed genes that were responsible for the various observed phenotypes, we measured global transcript levels using RNA-seq (Supplemental Table 1). Previously we found that ~11% of the genome was differentially expressed in a *B. dolosa fixLJ* deletion mutant.^21^ There were 205 genes that were significantly downregulated and 302 genes significantly upregulated (*q* value <0.05, at least 2-fold difference) in *B. dolosa* carrying evolved FixL variant W439S (Figure 3E) compared to *B. dolosa* carrying the ancestral FixL W439. We analyzed for Gene Ontology (GO) terms that were enriched among the genes that were significantly differentially expressed. Surprisingly, there were no GO terms that were enriched among genes that were upregulated in *B. dolosa* carrying the ancestral FixL W439. However, there were four GO terms that were enriched among genes that were differentially expressed in *B. dolosa* carrying the evolved variant (FixL W439S, Figure 3F). Three of these GO terms are associated with motility or chemotaxis, and the fourth is associated with signal transduction. A subset of the genes associated with motility and flagellar assembly that were differentially up-regulated in *B. dolosa* carrying the evolved variant (FixL W439S) are listed in Table 1. We confirmed the differential levels of *fliC* and *motA* transcripts in *B. dolosa* carrying the ancestral FixL variant (W439), evolved FixL variant (W439S), and empty vectors using qRT-PCR (Supplemental Figure 1).

**Table 1.**
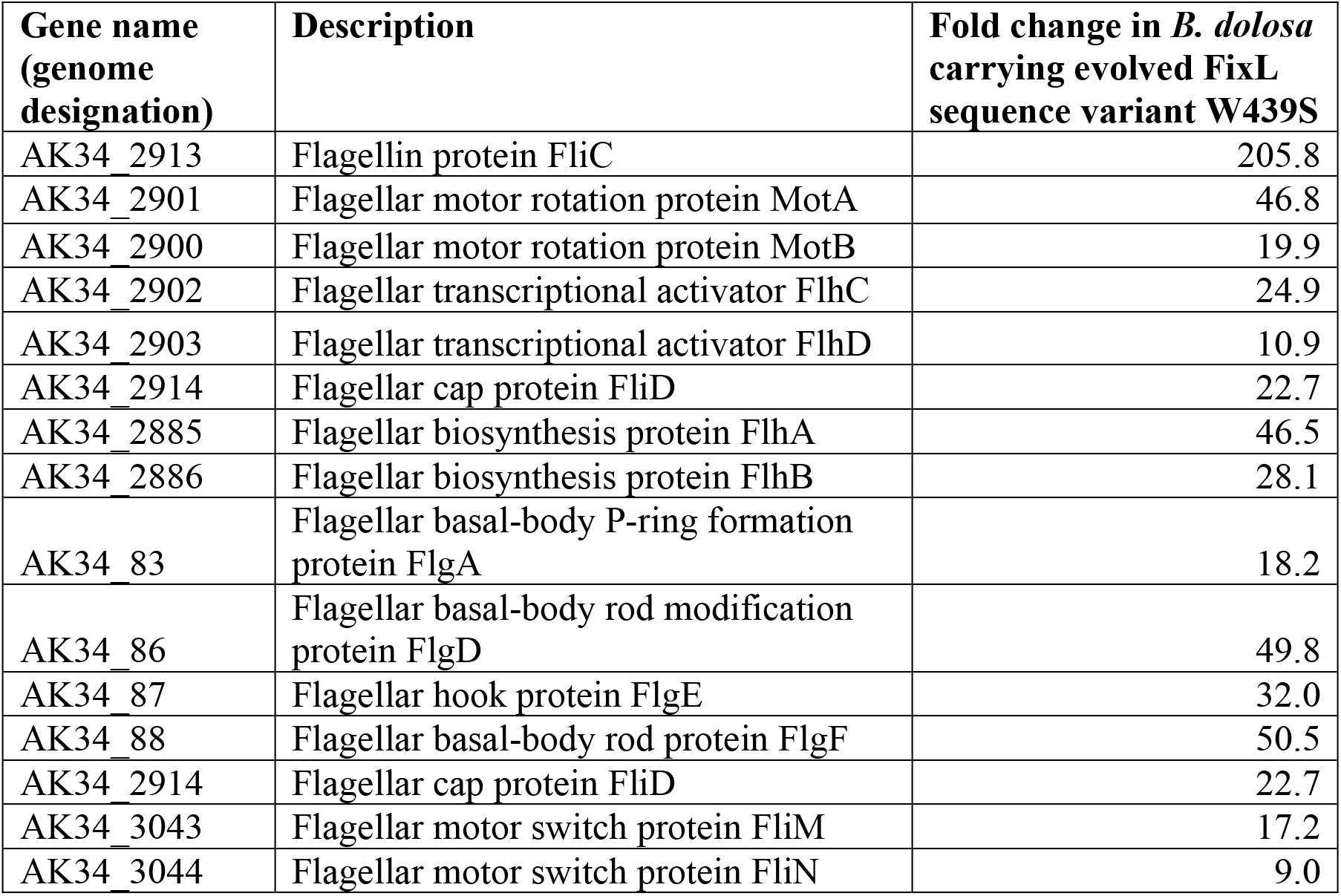
Selected genes associated with motility and/or flagellar assembly that are significantly differentially expressed in *B. dolosa* carrying evolved FixL sequence variant (W439S) relative *B. dolosa* carrying evolved FixL ancestral variant, W439.

| Gene name (genome designation) | Description | Fold change in *B. dolosa* carrying evolved FixL sequence variant W439S |
| --- | --- | --- |
| AK34_2913 | Flagellin protein FliC | 205.8 |
| AK34_2901 | Flagellar motor rotation protein MotA | 46.8 |
| AK34_2900 | Flagellar motor rotation protein MotB | 19.9 |
| AK34_2902 | Flagellar transcriptional activator FlhC | 24.9 |
| AK34_2903 | Flagellar transcriptional activator FlhD | 10.9 |
| AK34_2914 | Flagellar cap protein FliD | 22.7 |
| AK34_2885 | Flagellar biosynthesis protein FlhA | 46.5 |
| AK34_2886 | Flagellar biosynthesis protein FlhB | 28.1 |
| AK34_83 | Flagellar basal-body P-ring formation protein FlgA | 18.2 |
| AK34_86 | Flagellar basal-body rod modification protein FlgD | 49.8 |
| AK34_87 | Flagellar hook protein FlgE | 32.0 |
| AK34_88 | Flagellar basal-body rod protein FlgF | 50.5 |
| AK34_2914 | Flagellar cap protein FliD | 22.7 |
| AK34_3043 | Flagellar motor switch protein FliM | 17.2 |
| AK34_3044 | Flagellar motor switch protein FliN | 9.0 |

Among the genes that were differentially regulated we identified several differentially expressed genes that have homologs identified in other bacteria that play a role in c-di-GMP metabolism (Supplemental Table 2). This expression pattern suggests that there may be a decrease in intracellular c-di-GMP levels in the evolved variants and this may explain the increased motility and decreased biofilm seen in these constructs.^25–27^ Notably a homolog of the predicted phophodiesterase *cpdA* that could hydrolyze c-di-GMP, AK34_1958, was significantly downregulated in *B. dolosa* carrying the ancestral FixL variant (~8.5 fold, Table S2). *B. pseudomallei*^26^ and *B. cenocepacia* ^27,28^ mutants lacking *cpdA* had reduced motility consistent with the *B. dolosa* carrying the ancestral FixL variant which had lower *cpdA* transcript levels and lower motility. Similar to *B. pseudomallei* and *B. cenocepacia cpdA*, *B. dolosa* AU0158 *cpdA* has predicted GGDEF and EAL domains, but only the EAL domain is predicted to be enzymatically active based on amino acid sequence at the catalytic site. *B. pseudomallei*^26^ and *B. cenocepacia* ^28^ mutants lacking *cpdA* had increased levels c-di-GMP, but, surprisingly, there was a significant increase in intracellular c-di-GMP levels found in the *B. dolosa* construct carrying the evolved FixL variant (Supplemental Figure 2).

### Evolved FixL variants have different mechanisms of down-regulating fix pathway activity

Most TCS function by regulating gene expression when activated by a specific signal. The first component, the sensor kinase, senses the signal and autophosphorylates. The phosphate is then transferred to the second component, the response regulator, which can then regulate gene transcription.^20^ We conducted *in vitro* phosphorylation assays with recombinant FixL and FixJ proteins to measure the ability of the different FixL variants to phosphorylate themselves and then subsequently, FixJ. We hypothesized that FixL variants that had different phenotypes would have differing levels of autophosphorylation and/or phosphotransfer. Figure 4A shows that the ancestral FixL W439 had higher levels of autophosphorylation compared to the evolved FixL W439S. When the levels of autophosphorylation were quantified, the evolved FixL W439S had approximately 50% the level of autophosphorylation of the ancestral sequence (Figure 4B). Interestingly the other two evolved variants, G345S and R347H, also had similar levels of autophosphorylation compared to the ancestral variant (Figure 4A and 4B) despite making less biofilm and being more motile like the other evolved variant, W439S. Equal amounts of each of the purified protein were used in each reaction and each protein preparation was >90% pure (Supplemental Figure 3). When the autophosphorylation levels were quantified relative to the 1-minute time point, only the W439S autophosphorylation levels decreased while the other three variants increased or stayed the same (Supplemental Figure 4). We also measured the ability of the recombinant FixL variants to phosphorylate the response regulator FixJ. As the evolved variant W439S had lower levels of autophosphorylated FixL, that variant had lower levels of phosphotransfer to FixJ compared to the ancestral variant W439 (Figure 4C and 4D). When the level of phosphotransfer to FixJ was quantified relative to level of autophosphorylated FixL, the evolved variant W439S had a different profile compared to the ancestral variant and the other two evolved variants. The W439S had a more rapid phosphotransfer to FixJ and a greater level of dephosphorylation of FixJ (Figure 4D).

**Figure 4.**
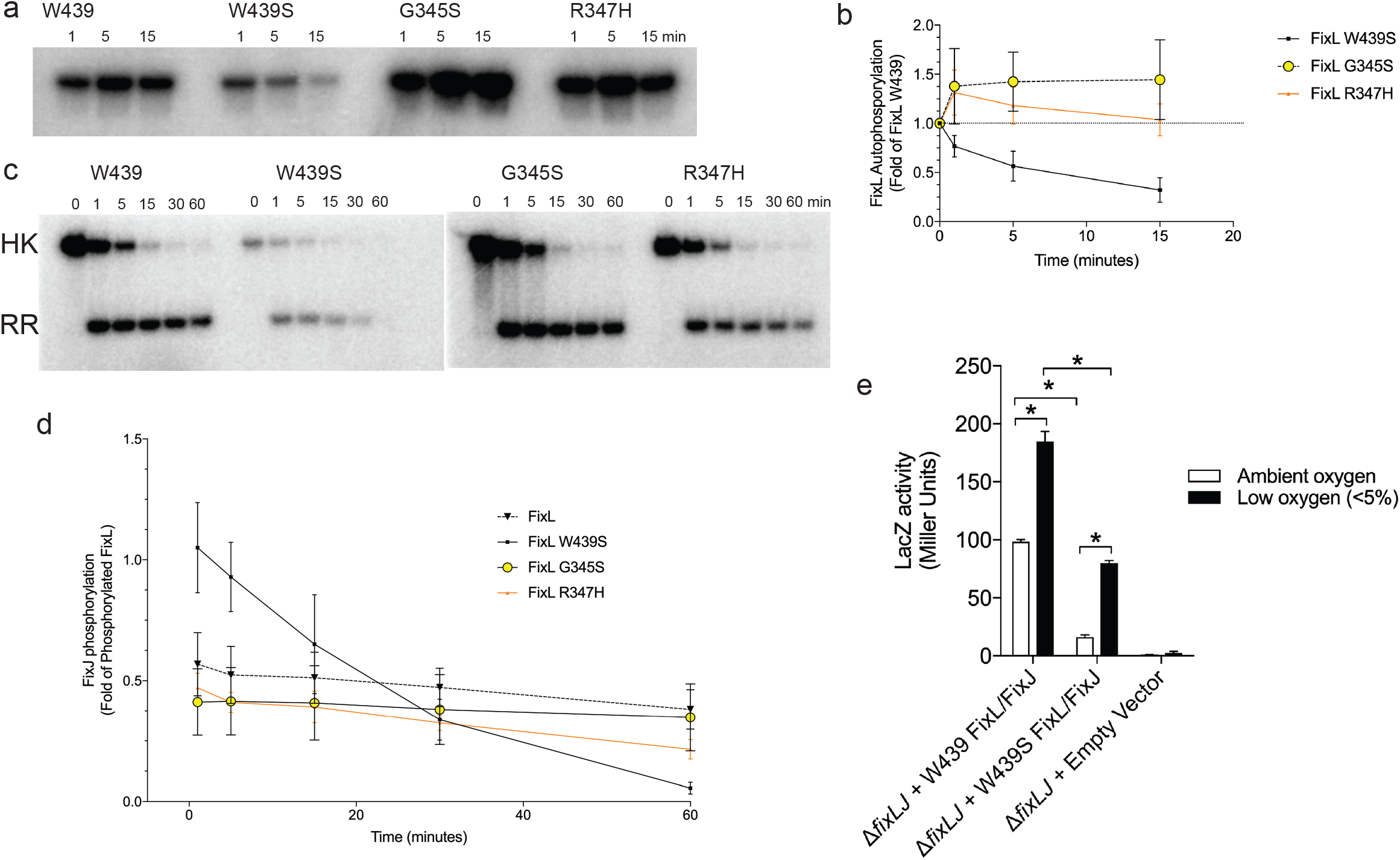
Evolved FixL variants have different mechanism of down regulating fix pathway phosphorylation. **(A)** Representative plots of autophosphorylation of *B. dolosa* FixL variants. **(B)** Density measurements from 3 independent experiments were normalized relative W439 at the same time point. *P<0.05 compared to W439 at the same time point by t test. **(C)** Representative plots of phospho-transfer to FixJ of *B. dolosa* FixL variants. **(D)** Density measurements from 3 independent experiments were normalized based on level the level of FixL phosphorylation at time 0 for each construct. *P<0.05 compared to W439 at the same time point by t test. **(E)** *B. dolosa* strain AU0158 constructs carrying FixL sequence variants or an empty vector carrying a p*fixK*-lacZ reporter plasmid21 grown in ambient or low (<5%) oxygen. Bars represent the means of triplicate biological replicates, and error bars represent one standard deviation (representative of three independent experiments). *P<0.05 by ANOVA with Tukey’s multiple comparison test.

Both of the other two evolved variants G345S and R347H had high initial levels of phosphotransfer (Figure 4C), but at later time points R347H had lower levels of phosphorylated FixJ compared to the ancestral FixL W439 suggesting R347H had increased levels of phosphatase activity. To confirm the downregulation of the *fix* pathway in an evolved variant, we measured *fix* activity using a *B*. *dolosa fixK* promoter-driven LacZ reporter plasmid using the pSCrhaB2 backbone.^21^ We found that both the ancestral (W439) and the evolved (W439S) FixL variants had an increase in *fix* pathway activity when the construct was grown in low oxygen (<5%), demonstrating that both variants are activated in oxygen tension (Figure 4E). Consistent with the *in vitro* phosphorylation experiments, the ancestral FixL (W439) had higher *fix* activity than the evolved variant (W439S) when grown in either ambient or low oxygen.

## Discussion

Previous work has determined that the BCC *fixLJ* system shows evidence of positive selective pressure during chronic lung infection in CF patients^17–19^ and is critical for BCC virulence.^21^ In this study we evaluated the function of evolved FixL sequence variations in both *B. dolosa* and *B. multivorans* by generating isogenic constructs carrying the evolved FixL variants. These evolved FixL variants were more virulent, more motile, and produced less biofilm. The FixL mutations that occurred during chronic infection downregulated the *fix* signaling cascade, demonstrating that high *fix* activity is detrimental for virulence (Figure 4). Interestingly, different FixL variants had different mechanisms of downregulating *fix* pathway activity, including decreases in autophosphorylation and increases in phosphatase activity.

We found that BCC carrying ancestral *fixL* sequences had reduced motility and reduced virulence in both murine and macrophage models compared to variants carrying evolved sequences (Figures 2 and 3) suggesting that the reduced motility was contributing to the reduced virulence, as has been suggested by other studies.^29–31^ Our previous work demonstrated that *B. dolosa* lacking flagella were equally virulent as parental, flagellated strains, suggesting motility plays a minimal role in the infection models evaluated.^21^ We analyzed the transcriptome of *B. dolosa* constructs carrying ancestral or evolved FixL variants to understand the mechanisms of the altered phenotypes seen between the two constructs. Among the genes that were identified to be differentially expressed between *B. dolosa* carrying different *fixL* sequences were genes that have homologs in other bacterial species that are involved in c-di-GMP metabolism (Supplemental Table 2).^25–27^ Surprisingly, *B. dolosa* carrying the ancestral FixL sequence that made increased levels of biofilm had lower levels of c-di-GMP than *B. dolosa* carrying the evolved FixL sequence variant (Figure S2). The increased biofilm seen in *B. dolosa* carrying the ancestral FixL sequence variant and in *B. dolosa* lacking *fixLJ* was independent of increased c-di-GMP levels, indicating that there are additional pathways involved in biofilm formation. Furthermore, high levels of c-di-GMP have been shown to stimulate the production of extracellular polysaccharides that leads to increased biofilm formation.^27,32^ Interestingly, the genes that are responsible for c-di-GMP induced production of polysaccharides (Bep, PNAG, and cepacian)^33–35^ were not differentially expressed by RNA-seq analysis (Supplemental Table 1) suggesting that c-di-GMP-independent mechanisms of biofilm production are involved.

One potential c-di-GMP-independent mechanism of biofilm production involves the *wsp* system that can promote biofilm formation in *B. cenocepacia* without direct activation of a diguanylate cyclase.^36,37^ Multiple components of the *wsp* system were 2-4-fold up-regulated in *B. dolosa* carrying ancestral FixL (Supplemental Table 2), suggesting a potential role of the *wsp* system in *fix* pathway-mediated biofilm formation. The contribution of biofilm formation to BCC virulence remains unclear, as constructs carrying ancestral FixL sequences produced <u>more</u> biofilm and were <u>less</u> pathogenic in our murine model of pneumonia than isogenic constructs carrying evolved FixL variants (Figures 2 and 3). Similarly in our previous study, *B. dolosa fixLJ* deletion mutants made more biofilm and were less pathogenic than the parental strain that contained *fixLJ*.^21^ These findings suggesting that BCC biofilms may not be beneficial for infection are supported by a study of CF lung explants from patients infected with *P. aeruginosa* and/or BCC and stained using species-specific antibodies.^38^ BCC bacteria were rarely found in biofilm-like structures while *P. aeruginosa* were often found in such structures, suggesting BCC do not form biofilms during infection. The conversion of BCC from a mucoid, biofilm-producing state to a non-mucoid phenotype is often observed during chronic infection and is correlated with worse clinical outcomes (notably different from *P. aeruginosa*, where mucoid strains are associated with clinical decline).^39^ Interestingly, these more virulent isolates are found early during infection for *P. aeruginosa* but are found later during infection for CF patients infected with BCC.^39,40^

Since increased biofilm formation is not associated with increased BCC virulence (Figures 2 & 3), we hypothesized that biofilm formation may allow for the *B. dolosa* to better survive within the soil. But the ability to make biofilms does not, on its own, confer the ability to survive within soil as *B. dolosa* lacking *fixLJ* was unable to persist in the soil at high levels (Figure 4) despite making higher levels of biofilm (Figure 3). The mutations in *fixLJ* that confer increased virulence are likely dead-end mutations as they make the bacteria less able to survive within the natural BCC reservoir. It is possible that these mutations make the bacteria more transmissible between human hosts -- future work will investigate this.

To better understand the mechanism of altered phenotypes seen in the constructs carrying different FixL variants we examined genes that had the largest magnitude of differential expression by RNA-seq. One such gene is an AraC family transcription regulator (AK34_4608) that was significantly downregulated (~95 fold) in *B. dolosa* carrying the evolved FixL variant (Supplemental Table 2). Expression of this gene is potentially detrimental to bacterial dissemination since a homolog of this gene was found to be downregulated in blood *B. cenocepacia* isolates taken from CF patients with cepacia syndrome compared to lung isolates taken at same time.^41^ Additionally, a homolog of this same gene was found to be downregulated in *B. pseudomallei* isolates from CF patients taken at late versus early time points.^42^ Another set of genes that were differentially expressed encode a putative CidA/CidB-like, holin/anti-holin system (AK34_3040, AK34_3041) that was significantly downregulated (~64- and 42-fold, respectively) in *B. dolosa* carrying the evolved FixL variant (Supplemental Table 2). Homologs of this system were found to up-regulated in *P. aeruginosa* when grown in vitro with *Staphylococcus aureus*,^43^ suggesting that upregulation of this system may result in more autolysis and increased biofilm formation.^44^ Further investigation is needed to determine the role of these genes in BCC virulence and their role in *fix* pathway-mediated phenotypes.

In this study, we have characterized the effects of mutations within the oxygen-sensing two-component system *fixLJ* system that arise during chronic infection in people with CF. Some of these mutations were associated with a period of clinical decline. Isogenic constructs carrying these evolved FixL sequence variants were more virulent, more motile, and produced less biofilm. Multiple studies have investigated the genetic and phenotypic changes that occur within *P. aeruginosa* and to a lesser extent BCC.^45–47^ During chronic infection, *P. aeruginosa* becomes less motile, produces more biofilm, shows increased antibiotic resistance, and has increased auxotrophy, ultimately leading to the evolution of reduced virulence of late isolates in animal models of infection.^45,46^ Less is known about phenotypic changes that occur during chronic BCC infection, and most prior work has focused on *B. cenocepacia*.^48^ The findings from this study demonstrate a novel way BCC adapt to the host by making the bacteria more pathogenic at the cost of being less able to survive within the environment.

## Methods

### Clinical data

Records of *B. dolosa-*infected patients were reviewed under Boston Children’s Hospital IRB protocol number M10-08-0370.

### Bacterial strains, plasmids, cell lines, and growth conditions

All strains used and generated in this study are listed in Table 2. *B. dolosa* strain AU0158 was obtained from Dr. John LiPuma (University of Michigan) and is an early isolate from the index patient from the *B. dolosa* outbreak (about 3 years into the outbreak). BCC and *E. coli* were grown on LB plates or in LB medium and supplemented with following additives: ampicillin (100 μg/mL), kanamycin (50 μg/mL for *E. coli,* 1 mg/mL for BCC), trimethoprim (100 μg/mL for *E. coli,* 1 mg/mL for BCC), gentamicin (15 or 50 μg/mL), chloramphenicol (20 μg/mL), or diaminopimelic acid (200 μg/mL). Plasmids that were used in this study are listed in Table 3. Human monocyte line THP-1 was obtained from ATCC and grown at 37°C with 5% CO_2_. THP-1 cells were cultured in RPMI-1640 medium containing 2 mM L-glutamine, 10 mM HEPES, 1 mM sodium pyruvate, 4500 mg/L glucose, and 1500 mg/L sodium bicarbonate, supplemented with 10% heat-inactivated FCS (Invitrogen) and 0.05 mM 2-mercaptoethanol. Low-oxygen environments were generated by the CampyGen Gas Generating System (Thermo-Fisher), and the low-oxygen concentration (<5%) is based on the manufacture specifications.

**Table 2.**
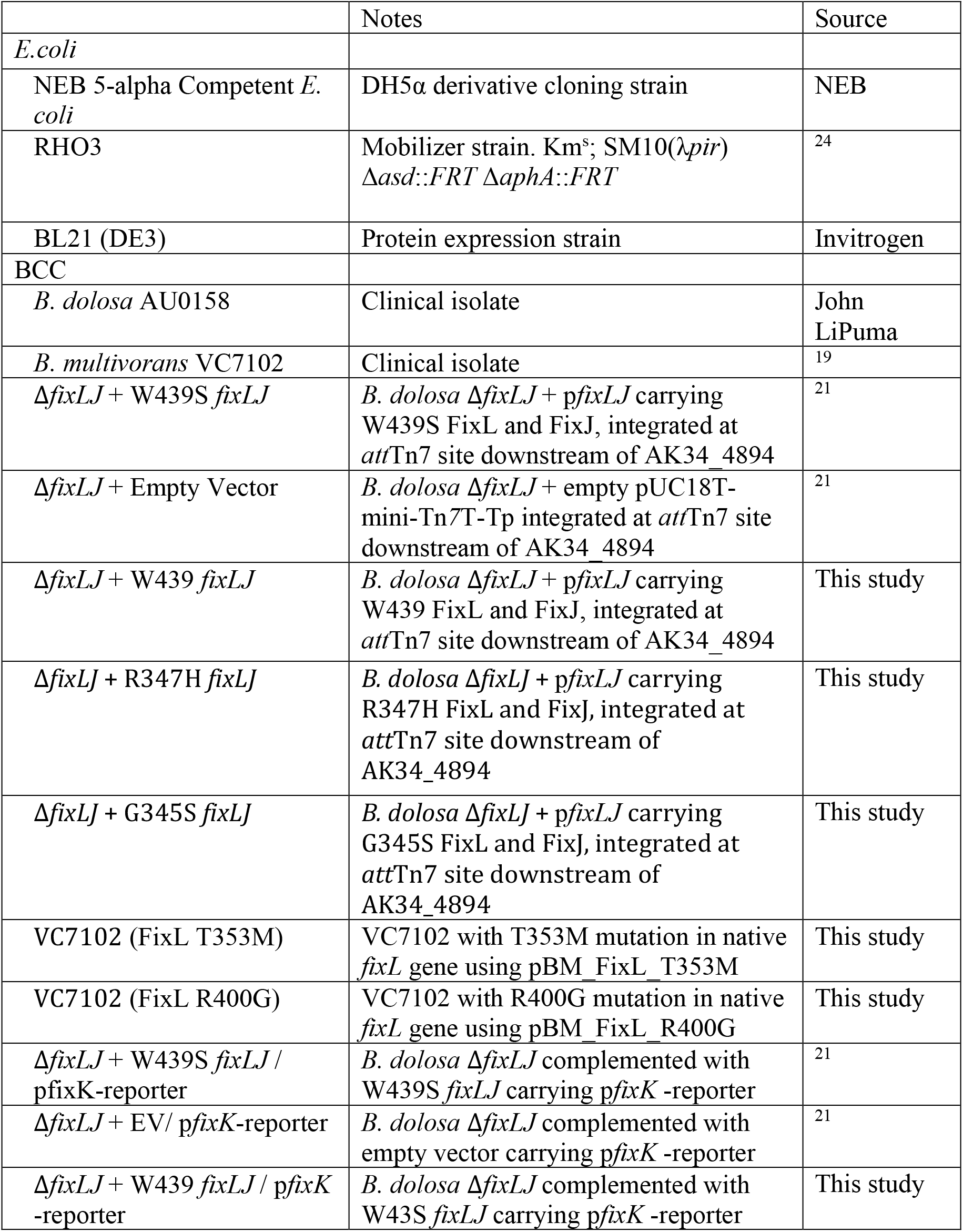
Strains used in this study.

**Table 3.**
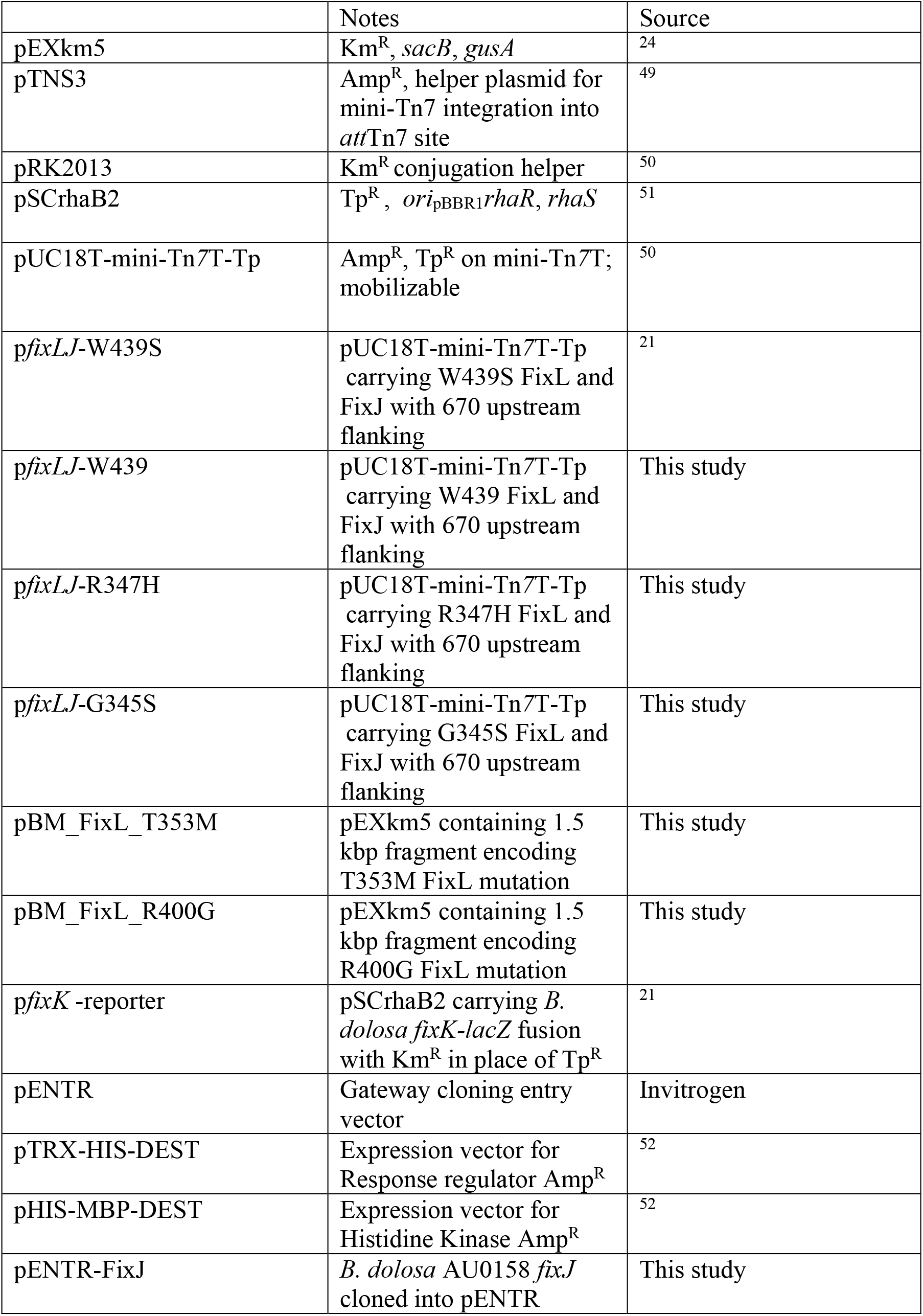
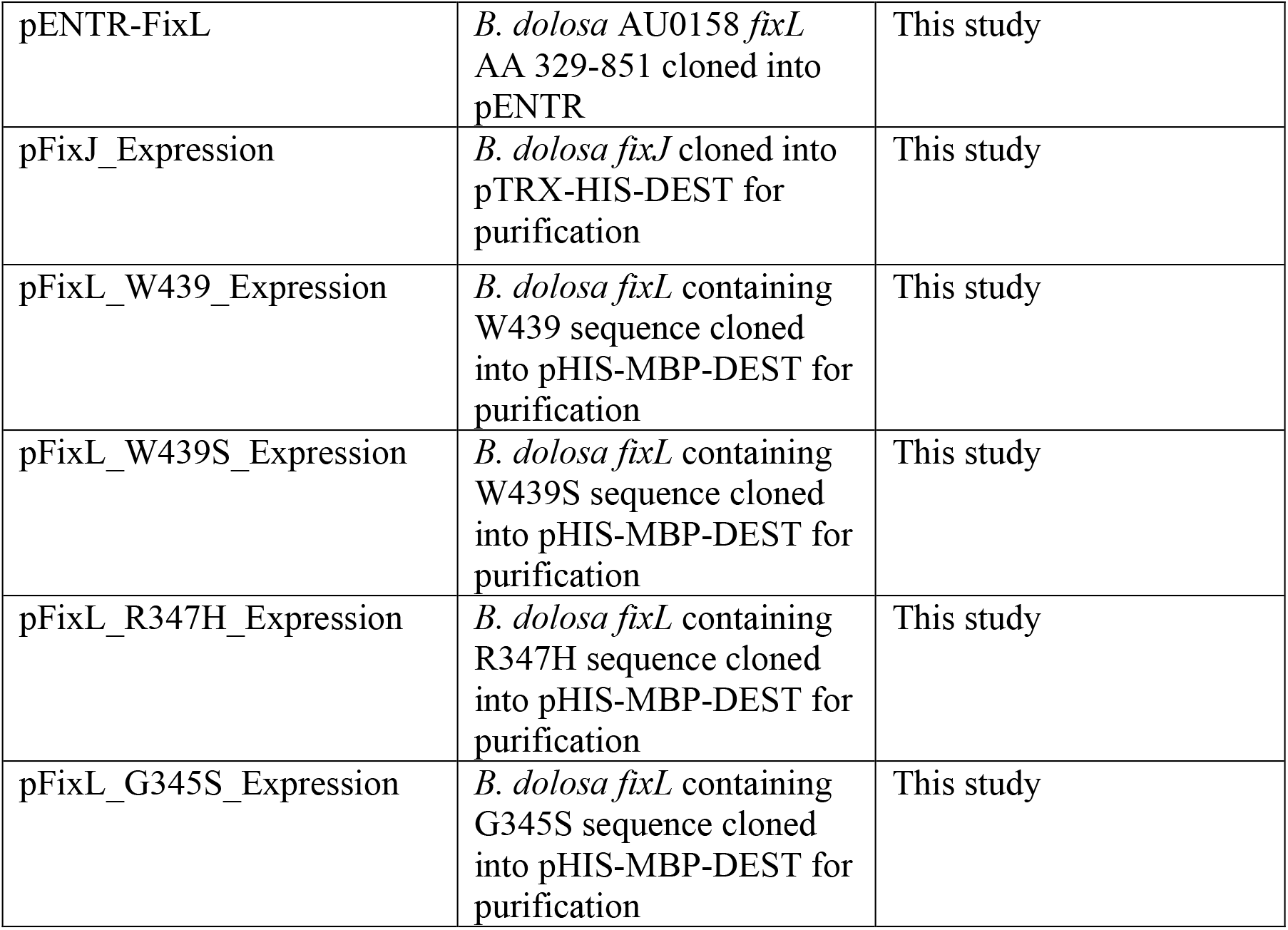
Plasmids used in this study.

|  | Notes | Source |
| --- | --- | --- |
| pEXkm5 | Km<sup>R</sup>, *sacB*, *gusA* | 24 |
| pTNS3 | Amp<sup>R</sup>, helper plasmid for mini-Tn7 integration into *attTn7* site | 49 |
| pRK2013 | Km<sup>R</sup> conjugation helper | 50 |
| pSCrhaB2 | Tp<sup>R</sup>, *ori*<sub>pBBR1</sub>*rhaR*, *rhaS* | 51 |
| pUC18T-mini-Tn7T-Tp | Amp<sup>R</sup>, Tp<sup>R</sup> on mini-Tn7T; mobilizable | 50 |
| *pfixLJ*-W439S | pUC18T-mini-Tn7T-Tp carrying W439S FixL and FixJ with 670 upstream flanking | 21 |
| *pfixLJ*-W439 | pUC18T-mini-Tn7T-Tp carrying W439 FixL and FixJ with 670 upstream flanking | This study |
| *pfixLJ*-R347H | pUC18T-mini-Tn7T-Tp carrying R347H FixL and FixJ with 670 upstream flanking | This study |
| *pfixLJ*-G345S | pUC18T-mini-Tn7T-Tp carrying G345S FixL and FixJ with 670 upstream flanking | This study |
| pBM_FixL_T353M | pEXkm5 containing 1.5 kbp fragment encoding T353M FixL mutation | This study |
| pBM_FixL_R400G | pEXkm5 containing 1.5 kbp fragment encoding R400G FixL mutation | This study |
| *pfixK* -reporter | pSCrhaB2 carrying *B. dolosa fixK-lacZ* fusion with Km<sup>R</sup> in place of Tp<sup>R</sup> | 21 |
| pENTR | Gateway cloning entry vector | Invitrogen |
| pTRX-HIS-DEST | Expression vector for Response regulator Amp<sup>R</sup> | 52 |
| pHIS-MBP-DEST | Expression vector for Histidine Kinase Amp<sup>R</sup> | 52 |
| pENTR-FixJ | *B. dolosa* AU0158 *fixJ* cloned into pENTR | This study |
| pENTR-FixL | *B. dolosa* AU0158 *fixL* AA 329-851 cloned into pENTR | This study |
| pFixJ_Expression | *B. dolosa fixJ* cloned into pTRX-HIS-DEST for purification | This study |
| pFixL_W439_Expression | *B. dolosa fixL* containing W439 sequence cloned into pHIS-MBP-DEST for purification | This study |
| pFixL_W439S_Expression | *B. dolosa fixL* containing W439S sequence cloned into pHIS-MBP-DEST for purification | This study |
| pFixL_R347H_Expression | *B. dolosa fixL* containing R347H sequence cloned into pHIS-MBP-DEST for purification | This study |
| pFixL_G345S_Expression | *B. dolosa fixL* containing G345S sequence cloned into pHIS-MBP-DEST for purification | This study |

### Genetic Manipulations and Strain Construction

To generate *B. dolosa* constructs carrying *fixL* mutations, we introduced the desired mutations using the Q5 Site-Directed Mutagenesis Kit (New England Biolabs) into p*fixK*.^21^ p*fixK* contains the AU0158 *fixLJ* sequence along with 670 bp upstream within the pUC18-mini-Tn7-Tp back bone allowing for stable chromosomally integration at an *att*Tn7 site.^23,50^ Point mutations were verified by Sanger sequencing. The *B. dolosa fixLJ* complementation vectors and the corresponding empty vector controls were conjugated into the AU0158 *fixLJ* deletion mutant with pRK2013 and pTNS3 using published procedures.^21^ Conjugants were selected for by plating on LB agar containing trimethoprim (1 mg/mL) and gentamicin (50 μg/mL). Insertions into the *att*Tn7 site downstream of AK34_4894 was confirmed by PCR. To generate *B. multivorans* VC7102 constructs carrying *fixL* mutations, we introduced the desired *fixL* mutations in the native *fixL* gene using the suicide plasmid pEXKm5.^24^ Briefly, approximately 1 kbp upstream and downstream of the desired mutation was PCR amplified, as was pEXKm5, and then the two fragments were joined using the NEBuilder HiFi DNA Assembly Master Mix (NEB) per manufacturer’s protocol. The plasmids were Sanger sequenced and transformed into RHO3 *E. coli* and then conjugated into *B. multivorans* VC7102.^23,49^ Conjugants were selected for on LB with kanamycin at 1 mg/mL. To resolve merodiploidy, conjugants were counterselected against by plating on LB with 15% w/v sucrose and incubated for 2 days at 30°C. Clones were screened for the introduction of desired mutation by PCR and Sanger sequencing.

Expression vectors for FixL and FixJ His_6_-tagged proteins were generated using the Gateway high-throughput recombinational cloning system (Invitrogen). The entire *fixJ* gene or *fixL* amino acids 329-851 (lacking transmembrane domains) were amplified from *B. dolosa* AU0158 and cloned in pENTR. Gateway LR clonase reactions were used to move *fixL* or *fixJ* into pHIS-MBP-DEST or pTRX-HIS-DEST, respectively, for expression.^52^ *fixL* mutations were introduced using Q5 Site-Directed Mutagenesis Kit (New England Biolabs) and sequences were confirmed by Sanger sequencing. To generate *fixK* reporter strains a *fixK-lacZ* fusion reporter was conjugated into *B. dolosa* constructs as previously described.^21^

### Bacterial invasion assays

The ability of *B. dolosa* to invade and persist within macrophages was determined using published protocols.^21^ Human THP-1 monocytes were differentiated into macrophages by seeding 1 mL into 24-well plates at 8×10^5^ cells/mL with 200 nM phorbol 12-myristate 13-acetate (PMA). THP-1 derived macrophages were infected with log-phase grown BCC washed in RPMI three times at ~2×10^6^ CFU/well (MOI of ~10:1). Plates were spun at 500 g for 5 minutes to synchronize infection and then incubated for 2 hours at 37°C with 5% CO_2_. To determine the total number of bacteria, wells were treated with 100 μL of 10% Triton-X100 lysis buffer (final concentration 1% Triton-X100), serially diluted, and plated to enumerate the number of bacteria. To determine the number of intracellular bacteria, separate infected wells were washed two times with PBS and then incubated with RPMI + 10% heat-inactivated FCS with kanamycin (1 mg/mL) for 2 hours. Monolayers were washed three times with PBS and lysed with 1% Triton-X100, serially diluted, and plated to enumerate the number of bacteria.

### Murine model of pneumonia

All animal protocols and procedures were approved by the Boston Children’s Hospital Institutional Animal Care and Use Committee (assurance number A3303-01). The specific protocol number is 18-01-3617R. All animal protocols are compliant with NIH Office of Laboratory Animal Welfare, Guide for the Care and Use of Laboratory Animals, The US Animal Welfare Act, and PHS Policy on Humane Care and Use of Laboratory Animals. Female C57BL/6 mice 6-8 weeks of age were obtained from Taconic Biosciences. Mice were maintained at the animal facilities at Boston Children’s Hospital. Mice were anesthetized with ketamine (100 mg/kg) and xylazine (13.3 mg/kg) given intraperitoneally. While the mice were held in dorsal recumbency, 10 μL of inoculum was instilled in each nostril (20 μL total). The inoculum consisted of log-phase *B. dolosa* washed in PBS and diluted to a concentration of ~2×10^10^ CFU/mL (4×10^8^ CFU/mouse). Mice were euthanized 7 days after infection by CO_2_ overdose, when lungs and spleen were aseptically removed. Lungs and spleens were weighed and placed into 1 mL1% proteose peptone in water, homogenized, and then serially diluted and plated on Oxidation/Fermentation-Polymyxin-Bacitracin-Lactose (OFPBL) plates.

### Biofilm formation

The ability to form biofilm on PVC plates was determined using published methods.^53^ Briefly, overnight cultures were diluted in Trypticase Soy Broth (TSB) with 1% glucose and pipetted into wells of a 96-well PVC plate. Plates were incubated 48 hours at 37°C, when unattached bacteria were washed with water. Biofilms were stained with 0.5% crystal violet and excess stain was washed away, and stain was solubilized with 33% acetic acid. The solution was transferred to a flat bottom plate, and then biofilm amount was quantified by measuring OD_540_.

### Motility assay

The ability of *B. dolosa* to swim was measured in low density LB agar using published methods.^54^ Briefly, 10 μL of overnight *B. dolosa* culture was plated in the center of low density (0.3% agar) LB plate. Plates were incubated agar side down for 48 hours at 37°C when swimming diameter was measured.

### RNA-seq

RNA was isolated from log-phase *B. dolosa* (two biological replicates per construct) using the Ribopure Bacterial RNA Purification Kit (Ambion) per manufacturer’s protocol, and contaminant DNA was removed using DNase. RNA was processed and libraries generated as previous published.^21^ Samples were sequenced using single end 50 bp reads using the Illumina HiSeq platform. Data analysis was done using Galaxy (usegalaxy.org).^55^ Reads (9-12 million reads per replicate) were trimmed using the Trimmomatic tool^56^ and mapped to the *B. dolosa* AU0158 genome (GenBank assembly accession GCA_000959505.1)^57^ using BowTie2 with very sensitive local preset settings.^58^ Differentially expressed genes were identified using CuffDiff using Benjamini–Hochberg procedure to determine the q value (*p* value corrected for multiple comparisons).^59^ Reads were deposited to BioProject PRJNA579568. GO terms that were enriched among genes that were differentially regulated were identified using GoSeq, using a Wallenius approximation and a Benjamini–Hochberg to determine a corrected p-value for multiple comparisons.^60^ GO terms that were considered to be enriched had and adjusted *p* value <0.05.

### qRT-PCR

cDNA was synthesized from 2 μg RNA using the ProtoScript II First Strand cDNA Synthesis Kit (NEB) per manufacture’s protocol. cDNA was cleaned using QIAquick PCR Purification Kit (Qiagen). Oligos to amplify *gyrB, rpoD,* and *fliC* were previously published,^21^ and *motA* was amplified using 5’-GTGAAGATCGGGCTCTTGT-3’ and 5’-GGACGTCTATATGGAGCTGATG-3’. Genes were amplified using oligos FastStart Essential DNA Green Master Mix (Roche) per manufacturer’s protocol. Expression was determined relative to *B. dolosa* AU0158 carrying the W439S FixL variant normalized by *gyrB* (AK34_3072) or *rpoD* (AK34_4533) expression using the ΔΔCt method.^61^ Both *gyrB* and *rpoD* had similar expression by RNA-seq between AU0158 and the *fixLJ* deletion mutant, and these genes have been used to normalize expression in *B. cenocepacia* in other studies.^62,63^

### Protein expression and purification

FixL and FixJ expression vectors were transformed into *E. coli* BL21 DE3 cells and protein was expressed and purified using nickel affinity columns following published protocols.^52^

### In vitro phosphorylation

Autophosphorylation and phosphotransfer assays were done as previously described.^64^ Briefly, FixL variants were used at a final concentration of 2.5 μM mixed with a final concentration of 5 mM MgCl_2_, 0.5 mM ATP, and 2.5 μCi of [γ^32^P]-ATP (stock of 6000 Ci/mmol 10 mCi/mL, from Perkin Elmer). Autophosphorylation reactions were performed at 30 °C, ambient oxygen, and stopped at the indicated time points by the addition of 4x sample buffer (200 mM Tris-Cl at pH 6.8, 400 mM DTT, 8% SDS, 0.4% bromophenol blue, 40% glycerol). For phosphotransfer assays, FixL variants were autophosphorylated using the above parameters at 30°C for 15 minutes and then incubated with reactions containing the response regulator FixJ and MgCl2 at final concentrations of 5 μM and 5 mM, respectively. Phosphotransfer reactions were run at 30 °C and ambient oxygen. Reactions were stopped at the indicated time points with the addition of 4x sample buffer. Samples were then run on an “any kD” BioRad mini-protean TGX gel for 50 minutes at 150 V. Gels were exposed to phosphor-screens for 4-5 hours so that phosphorylated protein bands could be observed. Screens were imaged using the Typhoon-FLA9500 imager with a “phosphor” setting and a resolution of 50 μm. Band intensity of phosphorylated proteins was quantified using ImageJ.

### Reporter assay

BCC carrying a *fixK-lacZ* report plasmid were grown overnight in LB with kanamycin (1 mg/mL). Cultures were subcultured in LB in ambient oxygen or LB that had been degassed in CampyGen Gas Generating System (Thermo-Fisher). Cultures were grown in ambient oxygen with shaking (200 rpm) at 37°C or within CampyGen Gas Generating System at 37°C for 4-6 hours. The level of *fix* pathway-driven LacZ activity was measured by determining Miller Units following published procedures.^65^

### c-di-GMP quantification

*B. dolosa* constructs were grown to stationary phase when 50 mL aliquots were spun down and c-di-GMP was extracted using ice-cold extraction buffer (methanol:acetonitrile:dH_2_O 40:40:20 + 0.1 N formic acid). c-di-GMP levels were measured using mass spectroscopy as previously described.^66^

### Data availability

RNA-seq reads have been deposited to BioProject PRJNA579568 https://www.ncbi.nlm.nih.gov/bioproject/579568.

## Acknowledgments

The authors would like acknowledge Brian Hsueh and Christopher Waters (Michigan State University) for measurement of c-di-GMP. This work was funded by the Cystic Fibrosis Foundation (PRIEBE13I0 to GPP) and the National Institutes of Health (R01GM110444 to VSC) and, in part, by the Richard A. and Susan F. Smith President’s Innovation Award (no number, to GPP) and by funds from the Translational Research for Infection Prevention in Pediatric Anesthesia and Critical Care (TRIPPACC) Program of the Department of Anesthesiology, Critical Care and Pain Medicine at Boston Children’s Hospital (no number, to GPP).

## Notes

https://www.ncbi.nlm.nih.gov/bioproject/579568

### Competing Interest Statement

The authors have declared no competing interest.

