## Supplemental Figures and Table S2 for "Host adaptations in the *fixLJ* pathway of the *Burkholderia cepacia* complex increase virulence"

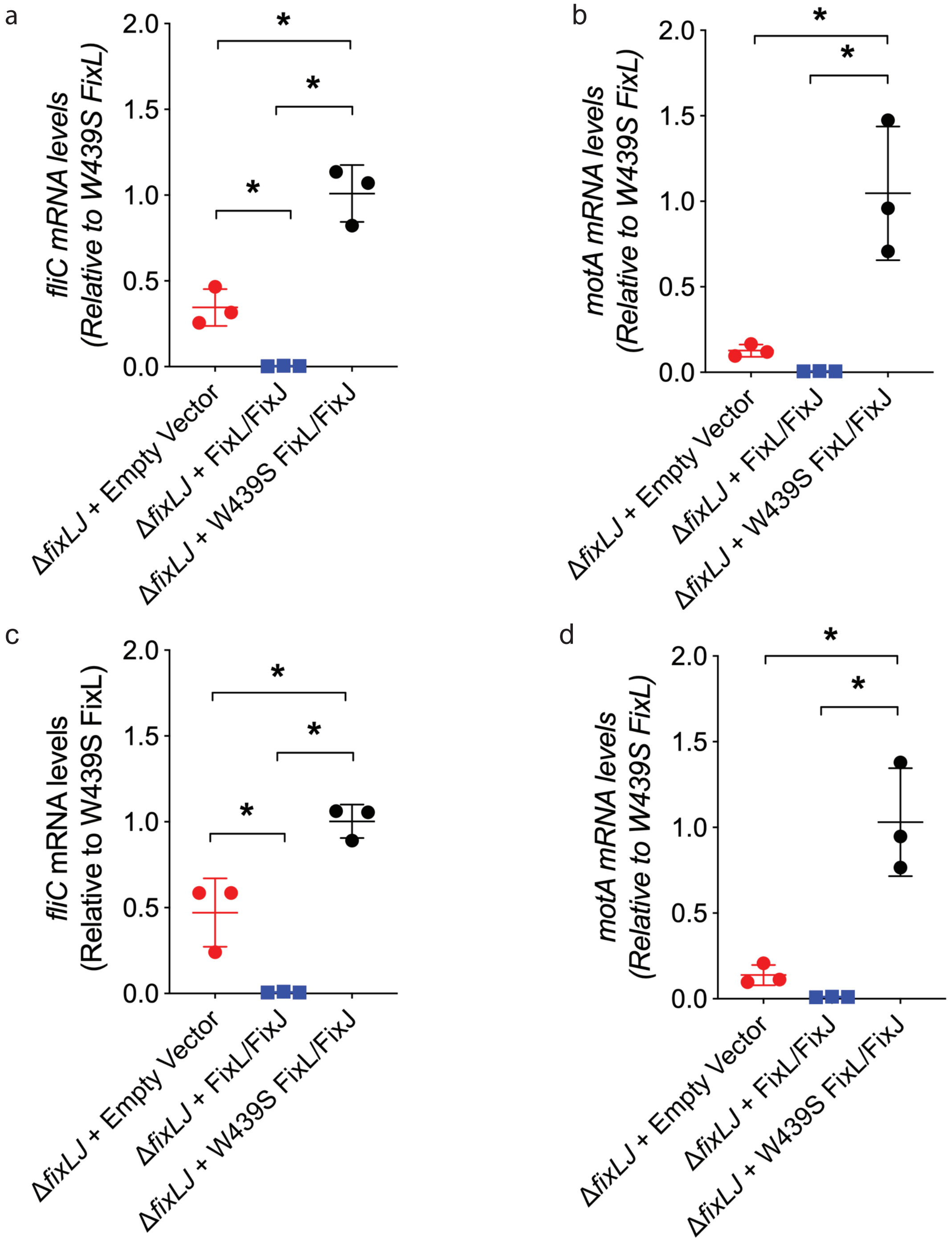


**Figure S1.** ***B. dolosa*** **carrying evolved FixL variant has higher levels of fliC and motA mRNA compared to** ***B. dolosa*** **carrying ancestral FixL variant.** Relative levels of *fliC* **(A&C)** and *motA* **(B&D)** in the *B. dolosa fixLJ* deletion mutant complemented ancestral FixL variant (W439), evolved FixL variant (W439S), or empty vector measured by qRT-PCR normalized to the levels of *rpoD* **(A-B)** or *gyrB* **(C-D)**. Plots are representative from 2 separate experiments with 2–3 biological replicates per experiment; error bars are S.D. *denotes P< 0.05 by ANOVA with Tukey's multiple comparison test.

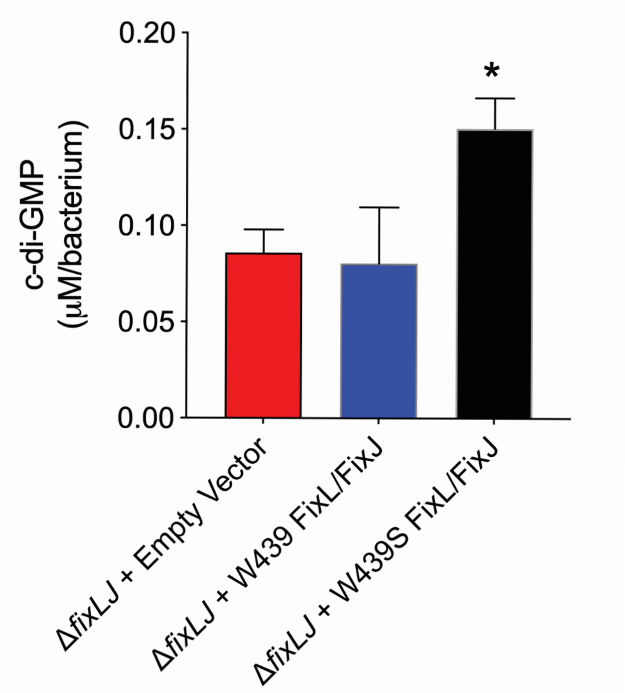


**Figure S2. *B. dolosa* carrying evolved FixL sequence variant have higher levels of c-di-GMP.** c-di-GMP was extracted from stationary growth *B. dolosa* and measured using mass-spectrometry. Mean data from 3 biological replicates and error bars are standard deviation. *P<0.05 by 1-way ANOVA with Tukey's multiple comparison test compared to Δ*fixLJ* +W439 FixLJ or Δ*fixLJ* + empty vector.

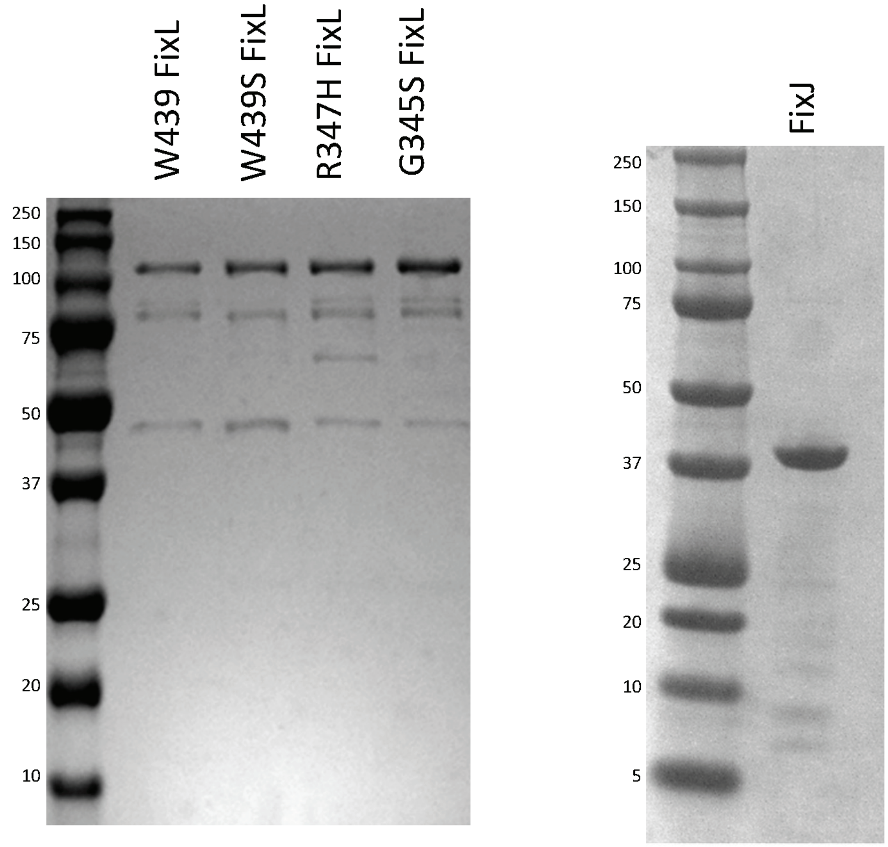


**Figure S3.** ***B. dolosa*** **FixL and FixJ protein preparations.** 1.2 µg of the various FixL sequence variants or 1 µg of FixJ was run on a SDS-PAGE gel and stained with Coomassie stain. Protein ladder sizes in kDa are listed. The expected size of the FixL is ~115 kDa and FixJ is ~40 kDa.

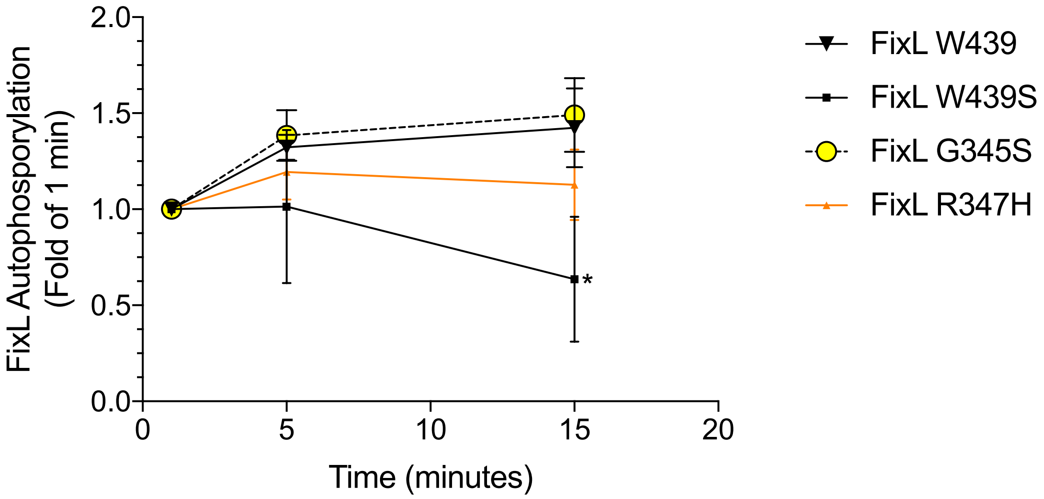


**Figure S4. W439S FixL variant has lower autophosphorylation activity than W439 (ancestral) variant.** Data from figure 4B normalized to 1 minute time point for each variant. *P <0.05 compared to W439 at time point by ANOVA with Tukey's multiple comparison test.

**Table S2.** Selected genes of interest that are significantly differentially expressed in *B. dolosa* carrying ancestral FixL sequence relative *B. dolosa* carrying evolved FixL sequence variant.

| **Category/ Gene name (genome designation)** | **Description** | **Fold change in *B. dolosa* carrying evolved FixL sequence** |
|---|---|---|
| c-di-GMP metabolism | | |
| AK34_137 | GGDEF and EAL domain containing protein | 3.5 |
| AK34_1958 | *cpdA* homolog<br>GGDEF and EAL domain containing protein | 8.5 |
| AK34_5467 | Response regulator containing an HD-GYP domain | 9.8 |
| *wsp* system | | |
| AK34_4625 | *wspA* | -2.9 |
| AK34_4626 | *wspHRR* | -4.0 |
| AK34_4623 | *wspC* | -3.0 |
| AK34_4621 | *wspE* | -3.0 |
| Others | | |
| AK34_4608 | AraC-like Transcriptional regulator | -94.6 |
| AK34_3040 | *cidA*<br>Holin-like protein | -63.9 |
| AK34_3041 | *cidB*<br>CidA-associated membrane protein | -41.6 |
